# Transparent Titanium Dioxide Nanotubes: Processing, Characterization, and Application in Establishing Cellular Response Mechanisms

**DOI:** 10.1101/330712

**Authors:** Jevin G. Meyerink, Divya Kota, Scott T. Wood, Grant A. Crawford

## Abstract

The therapeutic applications of titanium dioxide nanotubes (TiO_2_ NTs) as osteogenic surface treatments for titanium (Ti)-based implants are largely due to the finely tunable physical characteristics of these nanostructures. As these characteristics change, so does the cellular response, yet the exact mechanisms for this relationship remains largely undefined. We present a novel TiO_2_ NT imaging platform that is suitable for use with live-cell imaging techniques, thereby enabling, for the first time, dynamic investigation of those mechanisms. In this work, fabrication methods for producing transparent TiO_2_ NTs with diameters of 56 ± 6 nm, 75 ± 7 nm, 92 ± 9 nm, and 116 ± 10 nm are described. To demonstrate the diagnostic potential of these TiO_2_ NT imaging platforms, the focal adhesion protein vinculin and actin cytoskeletal filaments were fluorescently tagged in osteoblasts and real-time, high-resolution fluorescent microscopy of live-cell interactions with TiO_2_ NT substrates were observed. The scope of such a platform is expected to extend far beyond the current proof-of-concept, with great potential for addressing the dynamic response of cells interacting with nanostructured substrates.

## 1. Introduction

Titanium (Ti) and Ti-alloys have been used in orthopedic implants for several decades due to their high strength-to-weight ratio, excellent corrosion resistance, and good biocompatibility [1]. While Ti alloys boast impressive physical properties, limited early-stage osteointegration, owing to the bioinert natural oxide (TiO_2_) that forms spontaneously on Ti, can result in lengthy patient recovery times with no guarantee of implant success [2]. To improve bone-implant integration, the addition of macro-, micro-, and nano-scale features have proven to be an effective method for enhancing cellular function or improving mechanical fixation and ultimately reducing patient recovery time [3]. This has prompted researchers to decipher how these surface modifications influence cellular response, and what more could be done to improve the success of orthopedic implants [4]. Moreover, few of these surface modifications have been as successful when influencing cellular response as TiO_2_ nanotubes (NTs) [5-7]. Adapted from research efforts in dye-sensitized solar cells, a unique electrochemical fabrication method, capable of forming uniform porous alumina from aluminum (Al) substrates, was applied to Ti substrates to form the first documented TiO_2_ NTs [8-10]. More importantly, this low-cost electrochemical process can be applied to complex geometries, like Ti implants, to produce a uniform TiO_2_ NT coating with highly tunable physical characteristics [11-13]. Furthermore, the TiO_2_ NT coating is of the same chemical composition as the natural oxide that exists on all Ti materials. Today, the answer to exactly how TiO_2_ NTs influence cellular response mechanisms remains inconclusive, but has progressed to a point of general understanding where TiO_2_ NTs of a specific size (diameter) range can be used to elicit a desired cellular response [14, 15]. Interestingly, TiO_2_ NTs have been shown to both promote and discourage the clustering of focal adhesion proteins and integrins based on NT diameter, suggesting the physical characteristics of TiO_2_ NTs affects the absorption of adhesion proteins and that an optimal TiO_2_ NT diameter for protein absorption may exist [16]. Previous studies have investigated TiO_2_ NTs *in vitro*, confirming this ability to enhance bone cell adhesion, proliferation, and mineralization [13, 17-19], further stating that 15-20 nm TiO_2_ NTs supply an optimal substrate for increased mesenchymal stem cell adhesion and proliferation [13]. This work has shown that tube diameters exceeding 100 nm caused cells to experience significant elongation with relatively high cell mortality [16]. While these and like reports supply valuable insight describing the influence of TiO_2_ NT feature size on various cell types (e.g. mesenchymal stem cells [13, 16, 17, 20], endothelial [21], osteoblasts [12, 22-26], and osteoclasts [5, 14]), they fall short on definitively reporting the mechanism by which TiO2 NT diameters influence cellular response [15, 27]. Previous research has revealed that other factors like surface charge and material chemistry play critical roles in the adhesion of biological proteins and cellular morphology [16, 24, 28]. Furthermore, the enhanced adsorption of biological proteins to specific TiO_2_ NT diameters suggests that a proteins’ electric charge and relative size is sensitive to the unique negative charge and dimensions of each TiO_2_ NT [29]. The work of Kulkarni et al. looks extensively into the importance of nanostructure topography, where TiO_2_ NT diameter, convex/concavity of the inner and outer rim, overall length, intratubular spacing, and the innate negative charge have significant influences over the binding of positively and negatively charged proteins [30]. Work produced from this approach has begun to decipher the relationships between nanostructure, protein absorption, and the resulting cellular response mechanisms by investigating what TiO_2_ NT electro-physio-chemical properties are responsible for eliciting specific cellular responses using *in vitro* experiments and theoretical modeling [29, 31]. While these findings are crucial, discrepancies remain about how these properties play into the overarching influence TiO_2_ NTs have on cellular responses [6, 16, 20, 27].

Recent advances in live-cell microscopy techniques such as fluorescence-resonance energy transfer (FRET), fluorescence-lifetime imaging microscopy (FLIM), and lattice-light sheet microscopy (LLSM) have enabled accurate, high spatial and temporal imaging of molecular dynamic interactions within live-cells to observe elusive cellular functions [32]. FRET/FLIM microscopy has opened the doors to studying cell-substrate mechanisms, allowing researchers to age, track, and measure pico-Newton sensitive forces across the cell via FRET-tension sensors [33, 34]. In this work we present a novel, live-cell imaging platform that enables the use of fluorescence microscopy with 0.5-6.5 μm thick transparent TiO_2_ NT coatings adhered to coverglass to establish a method of capturing and quantifying these intricate live-cellular responses in real-time.

## 2. Materials and Methods

### 2.1 Opaque TiO_*2*_ Nanotube Coating Fabrication

Ti disks were cut from a commercially pure (cp)-Ti rod (99.7% pure, High-Strength Grade 5 Titanium, McMaster-Carr, Elmhurst, IL) and subsequently polished via metallographic grinding/polishing (MetaServ 250 and Vector LC 250, Buehler, Lake Bluff, IL). Specimens were initially ground using progressively finer (i.e. 400, 600, 800, and 1200 grit) silicon carbide grinding paper. Final polishing was carried out with 1.0 μm alumina powder (Pace Technologies, Tucson, AZ) slurry and Lecloth pad (LECO, Saint Joseph, MI), finishing with a colloidal silica/hydrogen peroxide solution and Chem 2 polishing pad (Pace Technologies, Tuscan, AZ). Before anodization, samples were consecutively sonicated in deionized water and methanol for 5 minutes each.

TiO_2_ nanotubes were prepared by anodic oxidation using an altered two electrode flat cell (Model K0235, Princeton Applied Research, Oak Ridge, TN). A platinum mesh cathode (Princeton Applied Research, Oak Ridge, TN) and Ti disk anode were connected to a DC power supply (Mo # E3612A, Agilent Technologies, Santa Clara, CA) while resistance was measured using KI-Tool software (Tektronix, Beaverton, OR) and a multimeter (Keithley 2100 Series: 6½-Digit USB Multimeter, Tektronix, Beaverton, OR). A constant working distance of 5.3 cm was used for all NH4F based electrolytes, with voltages of 20-60 V. Ammonium fluoride (≥99% Fisher Scientific, Pittsburg, PA) was dissolved in anhydrous ethylene glycol (EG) (anhydrous 99.8% Sigma Aldrich, St. Louis, MO) and deionized water while continuously stirring with a magnetic stir bar. Electrolyte compositions consisted of 0.55 wt.% ammonium fluoride (NH_4_F) and a range of 1-6 wt.% H_2_O in ethylene glycol. After anodization samples were rinsed with methanol and sonicated in methanol 30 min to remove nanopores. Once dry, samples were annealed (Thermolyne FB1415m, Thermo Scientific, Waltham, MA) for 1 h at 450 °C, and allowed to air cool. Before use, the samples were again rinsed with methanol and air dried.

### 2.2 Physical vapor deposition of Ti-on-glass

Physical vapor deposition (PVD) was employed to deposit a uniform thicknesses of Ti atop 25 mm diameter x 0.17 mm glass cover-slips (Cat. # 72225-01, Electron Microscopy Sciences, Hatfield, PA) serving as the transparent base. Glass targets were fixed to an aluminum plate that had been successively sonicated and rinsed with water, methanol, and acetone. Prior to PVD deposition the glass surface was plasma etched at 250 V for 30 min at a pressure of 1.0 ×;10^−5^ mbar with an argon flow of 145 sccm. During deposition, a 50 W bias was employed to the target, while a 375 V potential facilitated Ti deposition. Deposition thicknesses of ∼0.5, 1, 2, and 4 μm were produced by this method.

### 2.3 Transparent TiO_2_ Nanotube Coating Fabrication

Ti coated glass samples were first inspected and loaded into the custom electrochemical flat cell, exposing a 3.14 cm^2^ anodization area. Electrolyte was circulated via a peristaltic pump. A platinum mesh cathode and Ti coated glass anode were anodized using the same DC power supply, software, and multimeter stated in the previous NT fabrication section stated for opaque TiO_2_ NT fabrication. Constant anodization voltages ranged from 20-40 V, while 60 V anodization required a two-step anodization process (i.e. 60-40 V) to enable the formation of large diameter nanotubes while preventing premature consumption of the PVD Ti substrate. Here a constant potential of 60 V was applied until approximately half the available Ti was anodized. At this point, the potential was reduced (3 V/min) to 40 V and held constant until the transparent TiO_2_ film was produced and confirmed in real-time with current density values. Anodization times were carried out until transparent TiO_2_ NTs were generated from the deposited Ti substrate.

### 2.4 Coating Characterization

The surface of the NT coatings was observed using scanning electron microscopy (SEM, Zeiss Supra 40VP, Zeiss, Jena, Germany). Top-down images were generally taken at 30 kX and 50 kX magnification. AFM images were collected using a Bruker Nanoscale 5 AFM with ScanAsyst-Air software in multimode and a 2 nm radius pyramidal tip with 0.4 N/m spring constant (Bruker Nano Inc, Santa Barbara, CA). Characterization of coating/substrate cross-sections were carried out by carefully scratching the anodized Ti sample to examine the fractured and detached TiO_2_ coating.

The inner diameter (ID), outer diameter (OD), and NT lengths (or TiO_2_ coating thickness) were measured using SEM micrographs in conjunction with image analysis software (ImageJ, NIH, Bethesda, MD). Micrographs were captured from five different locations on each sample to quantify the degree of variability. Seventy-five measurements of NT inner diameter, outer diameter, and coating thickness were performed from three separate locations per sample. Substrate transparency (% of maximum intensity) was determined via a video spectral comparator (VSC6000/HS, Foster + Freeman Ltd. Vale Park, Evesham, Worcestershire, WR11 1TD United Kingdom).

### 2.5 Plasmid constructs, cell culture conditions and live-cell microscopy

DH5alpha strain of E. coli with GFP-mouse vinculin (Cat. #67935) and mCherry-UtrCH (Cat. #26740) bacterial stabs (plasmids) were purchased from Addgene, Cambridge, MA. Bacterial culture was conducted based on Addgene protocols and plasmid elution by miniprep kit (QIAprep Spin Miniprep Kit, Qiagen, Hilden, Germany). Vinculin was used to observe focal adhesion complexes at sites of cell-ECM interactions during proliferation and adhesion, whereas filamentous actin-bound calponin homology domains of utrophin were used to observe cell migration and morphology.

Pre-osteoblast mouse bone cells MC3T3-E1 subclone-4 (ATCC, Manassas, VA) were cultured in α-MEM media with 1% penicillin/streptomycin and 10% FBS. 150,000 cells were seeded on coverslips coated with fibronectin or TiO2 NT platforms. The cells were transfected with the above plasmids using lipofectamine 2000 (Fisher Scientific, Pittsburg, PA). Cells were imaged to observe focal adhesions and actin filaments using an Olympus IX-70 inverted microscope with a UplanFL N 100x/1.30NA oil objective. The focal adhesions were observed using a GFP filter cube and actin filaments using a tetramethylrhodamine-isothiocyanate (TRITC) cube. Images were captured by an iXon Ultra 897 EMCCD (Andor, Belfast BT12 7AL, United Kingdom) camera controlled by Micro-Manager Open Source Microscopy Software [35]. White-light live-cell images were carried out with a one-second acquisition period up to 45 mins after initial adhesion was observed. Real-time super-resolution radial fluctuations (SRRF) image capturing was carried out with 10 ms exposure times, EM gain of 250, a ring radius of 0.5, radiality magnification of 4, 6 axes in ring, and a three-second acquisition period using Andor’s iXon ‘SRRF-Stream’ functionality.

### 2.6 Statistical Analysis

All experiments were performed in duplicate or triplicate according to sample availability, and all quantitative data were presented as the mean (±) standard deviation. Statistical comparisons were made with one-way ANOVA and Tukey post-hoc analysis using GraphPad Prism v7.00 (GraphPad Software, La Jolla California USA).

## 3. Results and Discussion

### 3.1 TiO_*2*_ NT Fabrication Process Optimization

The anodic oxidation process was first optimized to enable fabrication of transparent TiO_2_ NT imaging platforms with controlled NT diameters. While the relationship between anodization conditions and nanotube dimensions are largely established [11, 36, 37], these works were not completed with transparency in mind. The primary challenge for transparent TiO_2_ NT production is to ensure the coating is fully developed prior to consumption of the thin PVD Ti substrate. It is well-known that increased anodization voltage increases NT diameter, however, increased voltage also increases the thickness and stability of a NP layer that rests on the surface of the NTs. Defining the electrochemical conditions necessary to produce TiO_2_ NTs free of NPs at each anodization voltage began with discerning electrolytic compositions effects on TiO_2_ NT and NP formation.

Figure 1 shows a top-down SEM image of a TiO_2_ NT coating before and after sonication to remove the NP layer. The NP layer is characterized by isolated pores (dark) encased by a TiO_2_ matrix (light), while the NT surface is characterized by pore-like openings surrounded by TiO_2_ walls and separated by voids. In the early stages of anodization, particularly during the application of high cell potentials (used to achieve large diameters), the NP layer is too dense and tightly adhered to be easily removed. With extended anodization duration, the NP layer will eventually succumb to field assisted chemical dissolution and be removed. When developing transparent TiO_2_ NTs, however, anodization time is limited by the amount of Ti substrate (1-4 μm thick PVD Ti) and the desired NT layer thickness to achieve transparency (thickness increases with anodization time).

**Figure 1:**
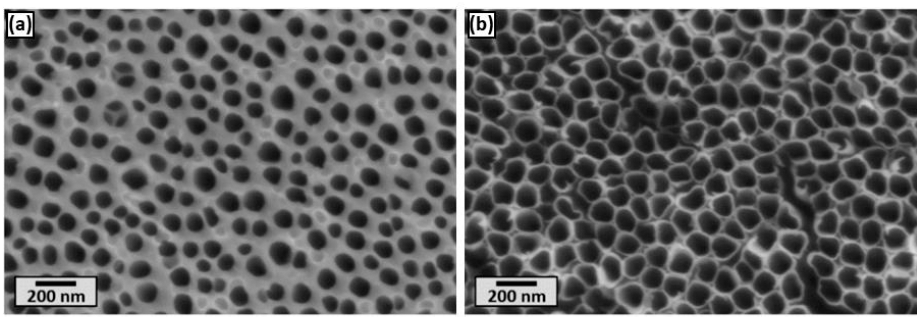
Top-down SEM micrograph of **(a)** TiO_2_ NPs, and **(b)** ultrasonically exposed TiO_2_ NTs. NTs are visible beneath the surface of the NPs in (a), illustrating that the NP and NT openings do not align.

To overcome this challenge, the influence of electrolyte water content on initial barrier layer thickness and associated TiO_2_ NP layer development was evaluated. The results of this evaluation (described below) demonstrated that an optimal water concentration of 2.5 vol% limits NP layer thickness while also preserving the quality of the NT surface appearance.

Top-down and cross-sectional SEM micrographs of TiO_2_ NT coatings fabricated using an electrolyte containing 0.14 M NH4F in ethylene glycol supplemented with varied amounts of deionized water (i.e., 1-6 vol% H_2_O) are shown in Figure 2, with corresponding TiO_2_ NT lengths (or coating thicknesses), measured by cross-sectional SEM analysis, shown in Figure 3. From inspection of Figure 2, increasing the electrolyte water concentration (above 3 vol %) resulted in an increasing presence of the NP layer after 30 min of anodization (Figure 2). It has been reported that minimizing water content results in a lower initial oxide (barrier) layer thicknesses, which directly translates to a reduction in NP layer thickness [38-40]. Because organic based electrolytes, such as ethylene glycol, contain limited O^−2^ concentration, even small changes in water concentration can have a significant impact on nanostructure formation and final morphology [38, 39, 41]. Increasing the ratio of F-to O^−2^ ions favors F^−^ assisted dissolution of TiO_2_, resulting in a thinner TiO_2_ barrier layer that is quickly penetrated to initiate TiO_2_ NT growth [42]. In this regard, at 1 vol% water content, the NT surface is clearly revealed (Figure 2a) without the presence of a NP layer. However, the NT layer surface has a mottled appearance owing to the lack of a protective barrier layer (due to low O^−2^ concentration), and associated NP layer, during the initial stages of NT growth. The continued chemical dissolution (due to high F^−^ concentration) of the surface of the NT layer throughout the anodization process also contributes to the mottled appearance. Continued etching of the surface of the NT layer and reduced oxygen content limits the NT growth rate, consequently limiting the thickness of the coating (Figure 3).

**Figure 2:**
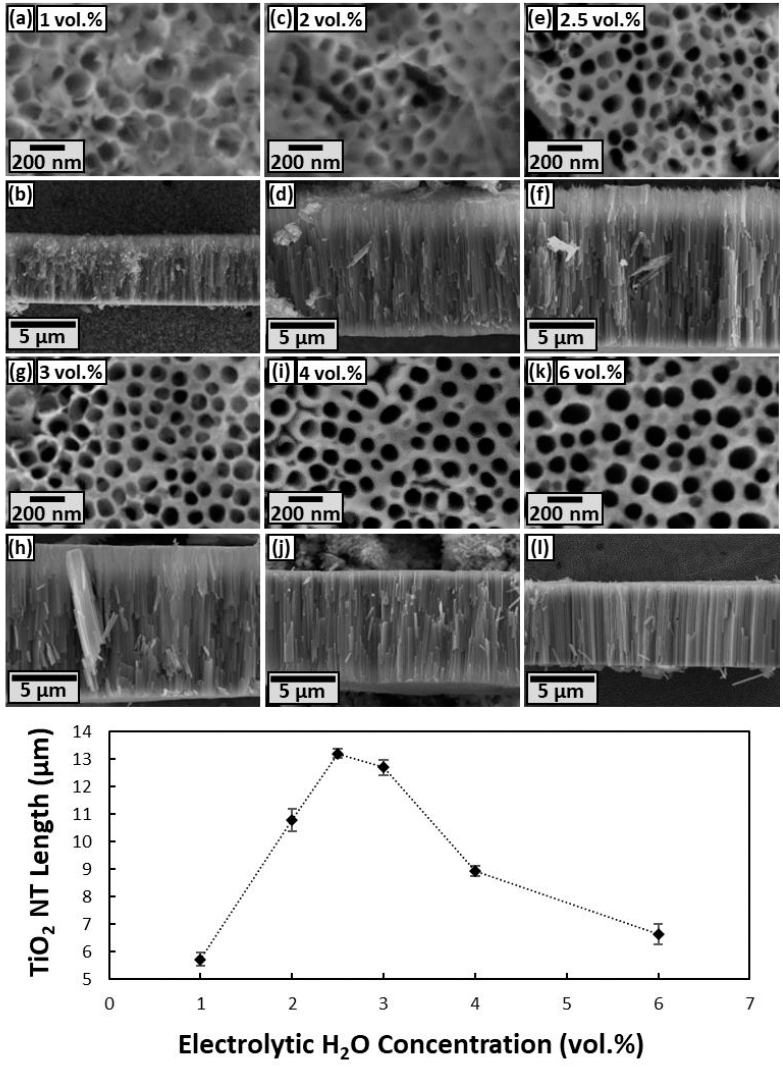
Top-down and cross section SEM micrographs of opaque TiO_2_ NTs/NPs anodized with **(a-b)** 1, **(c-d)** 2, **(e-f)** 2.5, **(g-h)** 3, **(i-j)** 4, **(k-l)** 6 vol.% H_2_O, 0.14 M NH_4_F, and ethylene glycol at 60 V for 30 mins.

**Figure 3:**
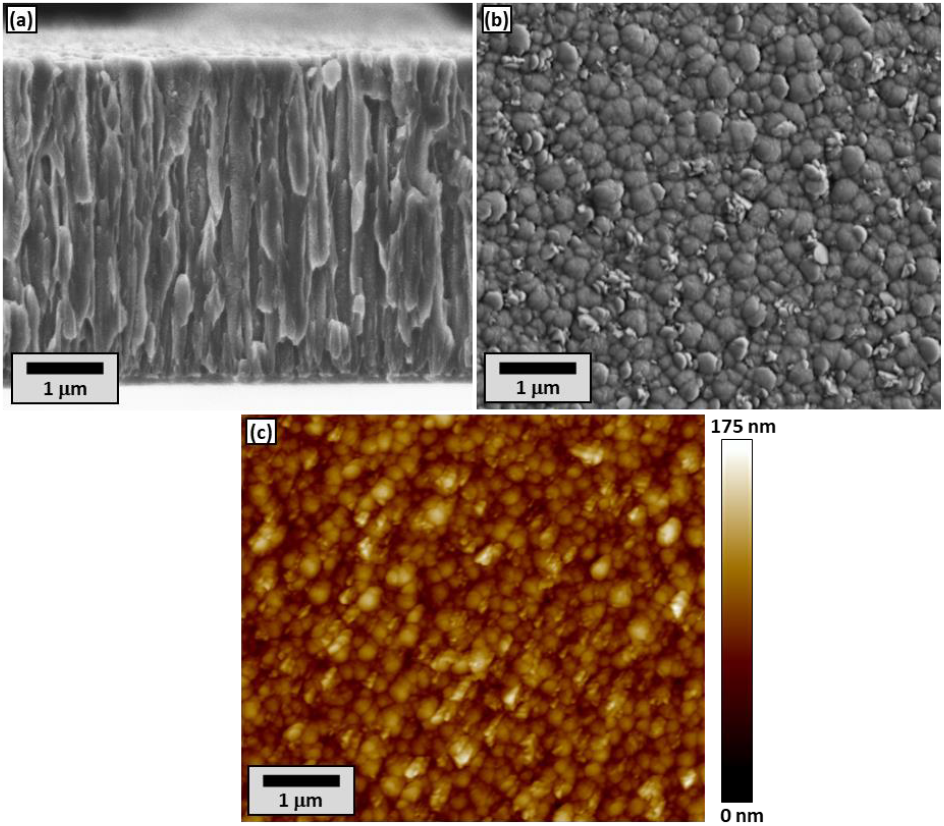
PVD Ti-on-glass visualization via **(a)** SEM cross-sectional, **(b)** SEM top-down, and **(c)** AFM top-down profile.

Increasing water concentration promotes oxide formation, resulting in the formation of a thick, dense NP layer during the initial stages of anodization that protects the underlying NT surface from chemical dissolution, and also permits rapid growth of the NT layer [43]. In addition, higher O^−2^/F^−1^ ratios favor increased oxidation and dissolution at the substrate/NT interface, which also increases the NT growth rate. As such, increasing the water content results in an increase in NT layer thickness (Figure 2). However, as the water content increases beyond 2.5 vol%, the increased thickness of the barrier layer that forms in the initial stages of anodization begins to limit O^−2^ and F^−^ ion transport to the Ti substrate which severely limits NT growth rate. Consequently, as water content increases above 2.5%, the NT thickness decreases with increasing water content (Figure 2). Therefore, based on these results an optimal water concentration of 2.5 vol% was identified that limits NP layer thickness while also preserving the quality of the NT surface appearance. Moreover, the thin NP layer present on the surface (Figure 2e) was easily removed via sonication in methanol to reveal the underlying TiO_2_ NTs, as shown in Figure 1b.

### 3.2 Transparent TiO_2_ NT Coating Fabrication

To fabricate transparent TiO_2_ NT coatings, a thin layer of Ti was deposited on glass substrates and subsequently subjected to electrochemical processing to form TiO_2_ NTs. Figure 3 shows a cross-sectional SEM micrograph and a top-down AFM surface profile of a PVD Ti coating on a glass substrate prior to anodization. From inspection of Figure 3a, the Ti coating is characterized by a columnar structure with a uniform thickness. In addition, the surface of the coating has a nanoscale surface texture with a surface roughness (Ra) of ∼3.9 nm. This structure is common in metallic PVD coatings [44]. The Ti coating thickness was controlled by deposition time and was adjusted to produce coatings with Ti thicknesses of 0.5, 1, 2, or 4 μm.

To demonstrate the overall transparent TiO_2_ NT coating fabrication process (Figure 4), a transparent glass coverslip was first imaged while resting on top of the logo of the South Dakota School of Mines & Technology printed on paper (Figure 4a). A glass coverslip was then placed on top of the same logo and imaged following PVD deposition of a 4-μm-thick Ti coating (Figure 4b). Finally, the Ti layer was subjected to anodic oxidation at an applied potential of 40 V, producing a transparent TiO_2_ NT coating (Figure 4c-e) through which the logo on the paper beneath is clearly visible (Figure 4c). The final thickness of the transparent NT coating in Figure 4e was approximately 7 μm. The development of this low-density oxide nanostructure resulted in a final TiO2 NT coating thickness that is much greater than the initial Ti layer thickness.

**Figure 4:**
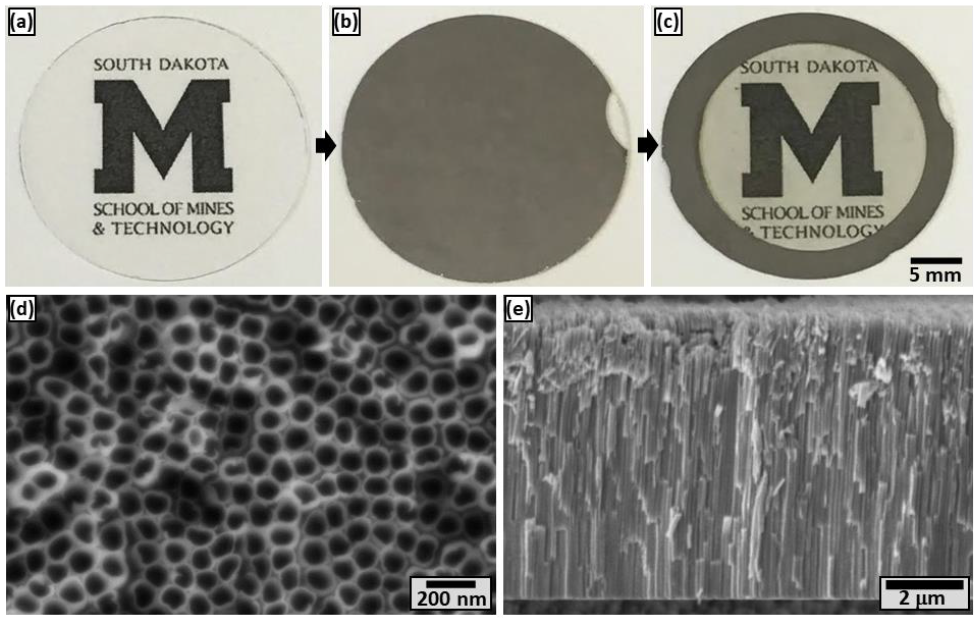
Visual representation of the overall fabrication method used to produce transparent TiO2 NTs, starting with photographs of a South Dakota School of Mines & Technology logo beneath a glass substrate **(a)** in as-received condition, **(b)** after PVD coating with a 4-μm-thick Ti coating, and **(c)** after anodizing at 40 V to form transparent TiO_2_ NTs. SEM **(d)** top-down and **(e)** cross-sectional micrographs of the transparent coating shown in (c) confirm the presence of TiO_2_ NTs.

To characterize the relationship between transparent TiO_2_ NT diameter and voltage, TiO_2_ NT coatings were produced with varying applied potentials (20-60 V) and a starting Ti thickness of 2 μm (Figure 5). Both the inner and outer diameters of NTs were found to increase with increasing anodization potential (Figures 5), which is in line with previous studies [45, 46]. Increasing the anodization potential increased the wall thickness of NTs (measured as the difference between OD and ID), as well.

**Figure 5:**
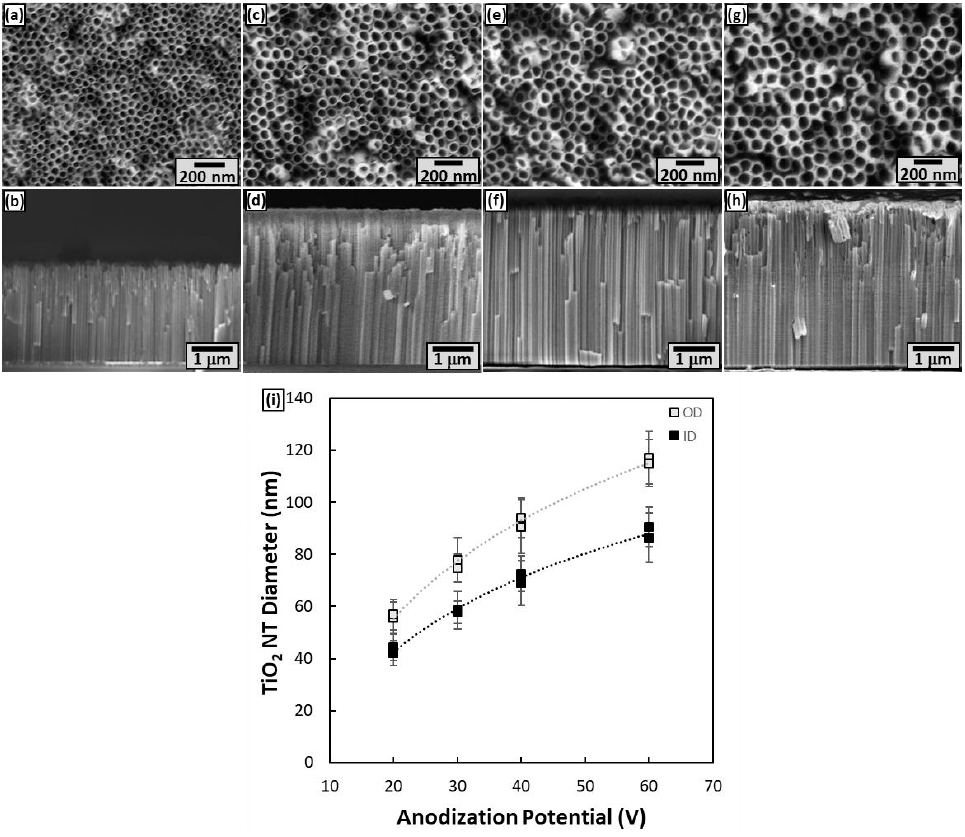
Top-down and cross section SEM micrographs of transparent TiO_2_ NTs anodized at **(a-b)** 20 V, **(c-d)** 30 V, **(e-f)** 40 V, and **(g-h)** 60-40 V with **(i)** associated inner diameters (ID) and outer diameters (OD) 56±6 nm, 75±7 nm, 92±9 nm, and 116±10 nm.

**Figure 6:**
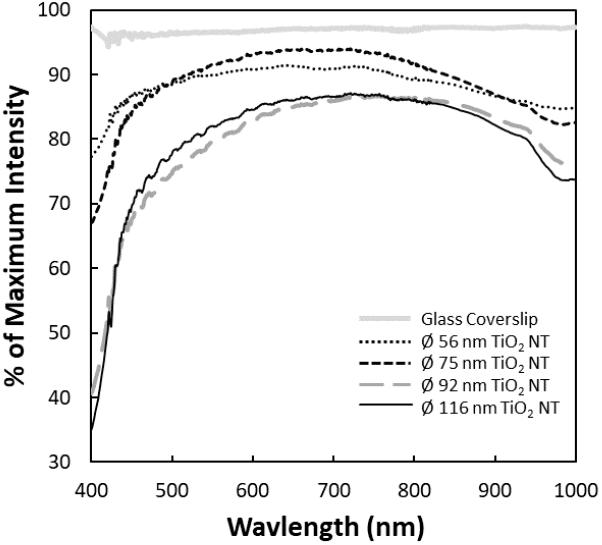
Transparency (% of maximum intensity) of each TiO_2_ NT coating fabricated from a 2 μm Ti substrate anodized to produce outer diameters (OD) of 56±6 nm, 75±7 nm, 92±9 nm, and 116±10 nm.

When fabricating TiO_2_ NTs with an applied potential of 60 V, enhanced TiO_2_ NTs growth rates resulted in rapid consumption of the 2 μm Ti substrate. This rapid growth limits chemical dissolution of the NP layer leaving an intact NP layer after anodization that cannot be removed with routine sonication. To circumvent this issue, a two-stage anodization, outlined in Section 2.3, was applied to reduce the growth rate and prolong anodization, permitting sufficient chemical dissolution for removal of the weakened NP layer. Thus, adjusting the applied potential throughout the anodization process allowed for precise control over the TiO_2_ NT growth rate and yielded fully transparent large diameter TiO_2_ NTs, without the presence of a NP layer, despite the limited initial Ti substrate thickness. It is important to note that the NT diameter present on the specimen surface is fixed by the initial anodization potential and unaffected by the two-stage process.

TiO_2_ NTs were obtained with outer diameters of 56±6 nm, 75±7 nm, 92±9 nm, and 116±10 nm for 20, 30, 40, and 60-40 V specimens, respectively (Figure 5). Interestingly, Ti substrates with initial thickness of 2 μm resulted in TiO2 NTs approximately 3.8 μm in length when anodized at potentials greater than 30 V (Figure 5c-h). The 20 V specimen, on the other hand, resulted in significantly shorter NTs. It is well-known that the overall growth rate of TiO_2_ NTs directly corresponds to applied potential, meaning longer durations are required to oxidize the same thickness of Ti when a lower potential is applied. As this anodization time increases, the NP layer and underlying NTs experience prolonged periods of chemical dissolution, effectively shortening the TiO_2_ NTs once TiO_2_ NPs are consumed. Thus, the 20 V specimen has a lower thickness because of the prolonged duration of anodization and associated chemical dissolution that occurred at the top of the NT coating while the 30, 40, and 60-40 V specimens have equivalent thicknesses that are fixed by the total amount of Ti available for anodization (i.e., 2-μm-thick PVD layer).

Thus far, we have demonstrated our ability to control NT dimensions while producing transparent TiO_2_ NT coatings. This is an important development, as it will permit the evaluation of the influence of nanostructures on biological responses using this platform. However, to be an effective tool for these evaluations, it is also critical to evaluate the influence of variations in nanostructure and coating thickness on optical transparency.

Transmission efficiency was measured at TiO_2_ NT lengths of 633±47 nm to 6.5±0.4 μm at 450-650 nm as shown in Figure 7. These wavelengths were selected as they are common excitation/emission ranges for fluorescent proteins used in live-cell imaging. TiO_2_ NTs < 5 μm long yielded highly transparent substrates >75%. Transparencies of 55-70% were observed with TiO_2_ NT 6.5±0.4 μm in length, while all reported fabrication conditions produce transparent TiO_2_ NT platforms that exhibited decreasing transparency with increasing TiO2 NT length.

**Figure 7:**
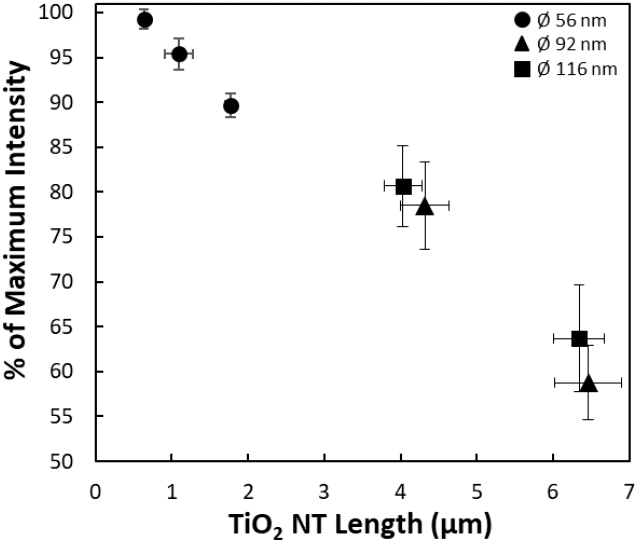
Average percent of maximum transmitted intensity of 450-650 nm wavelengths through transparent TiO_*2*_ NTs of 56, 92, and 116 nm OD, anodized from 0.5-4 μm Ti films atop glass coverslips to generate the TiO_*2*_ NTs lengths (*P*<0.0001 for all conditions).

### 3.3 Live-cell microscopy with Transparent TiO_2_ NT Coatings

To demonstrate functionality of the transparent TiO_2_ NT platform for use with live-cell imaging, epifluorescence images of live pre-osteoblast cells adhered to NTs with lengths of 6.5 ± 0.4 μm were obtained. Even with the lowest observed transparency of ∼55%, 6.5 ± 0.4 μm long TiO_2_ NTs still performed exceptionally well during live-cell imaging trials (Figure 8). The TiO_2_ NT platforms used in Figure 8 were comprised of the longest TiO_2_ NTs produced by the methods outlined in Section 2.3, yet permitted high-clarity imaging. From this, we infer that shorter TiO_2_ NTs with higher optical clarity can also be used in this manner to produce the same or better-quality images. It is well understood that as time progresses, so does the outgrowth of adherent cells, with clustering of vinculin apparent along actin filaments. The cells captured in Figure 8 exhibit the influence that different substrates (e.g., fibronectin-coated glass (Figure 8a-c) and TiO2 NTs (Figure 8d-f)) can have on cell morphology.

**Figure 8:**
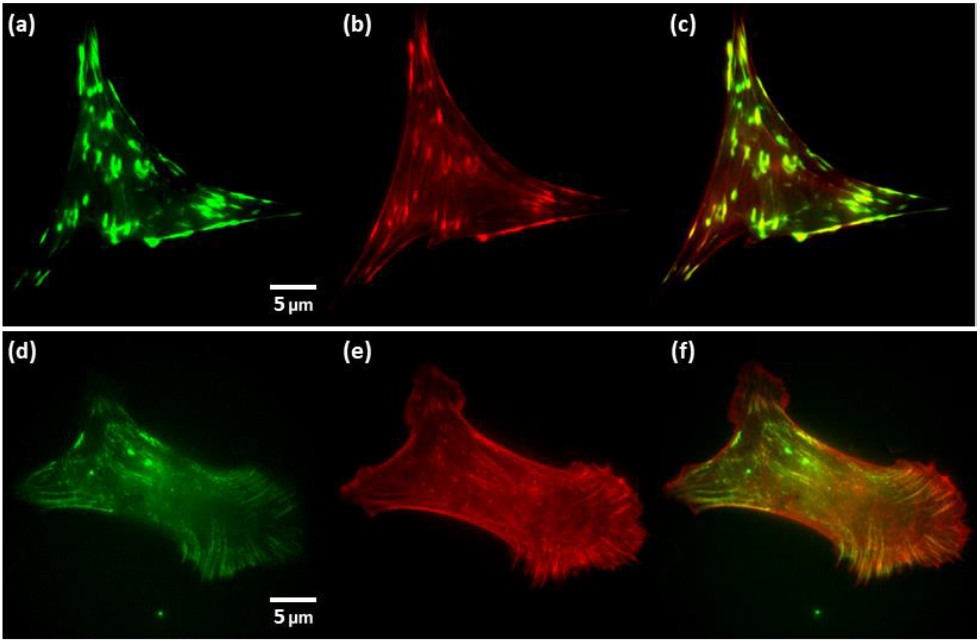
Live-cell images of two cells expressing both **(a, d)** green fluorescent vinculin and **(b, e)** red fluorescent actin seeded upon **(a-c)** fibronectin coated glass and **(d-f)** transparent 6.5±0.4 μm long TiO_2_ NT platforms, with **(c, f)** vinculin and actin images overlaid. Acquisition and display settings were identical between (a-c) and (d-f).

To further test the TiO_2_ NT platform functionality for time-lapse purposes, the initial contact and process of adhesion to the TiO_2_ NT substrate was targeted for imaging. To establish an effective time-scale imaging technique, white-light was first used on an inverted microscope frame to effectively capture this process and develop a timeline for adhesion, seen in Figure 9 and Supplemental Video 1. Once a cell was identified to have made initial contact with the substrate, live-cell time-lapse imaging was promptly initiated. The cell shown in Figure 9 shows clear signs of adhesion at 15 mins after initial substrate contact and spreading after 30 mins. The transparent nature of the NT platforms allowed for facile imaging of live cells interacting with NTs over time, thereby facilitating the ability to track and quantify cell growth in real time in future studies.

**Figure 9:**
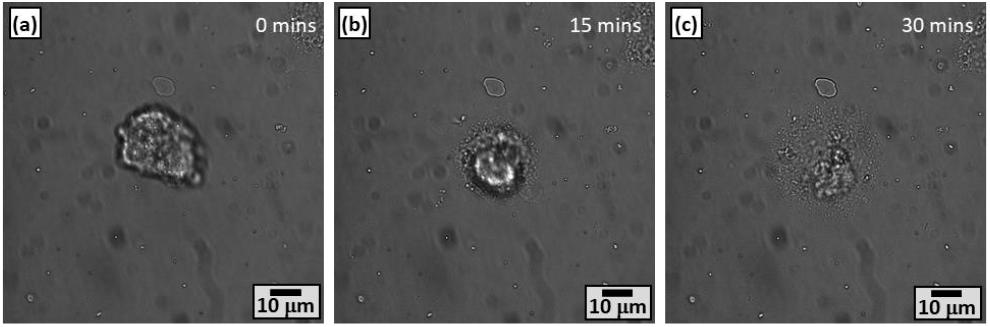
Images of a live-cell attaching to 56 ± 6 nm diameter, 633 ± 47 nm long, transparent TiO2 NTs at **(a)** initial adhesion, **(b)** 15 mins, and **(c)** 30 mins after contact.

Cells transfected with fluorescently-tagged actin filaments provided a sufficient model for demonstrating our platform’s functionality for live-cell fluorescence microscopy, as seen in Figure 10 and Supplemental Video 2. Time-lapse imaging of these transfected cells using an advanced fluorescence microscopy technique, super-resolution radial fluctuations (SRRF), provides clear visualization of cytoskeletal components and cell movement. Based on the actin movement, the cell in Figure 10 and Supplemental Video 2 appears to be dynamically probing the TiO_2_ NT substrate using filopodia and lamellipodia at 20-24 mins post-seeding.

**Figure 10:**
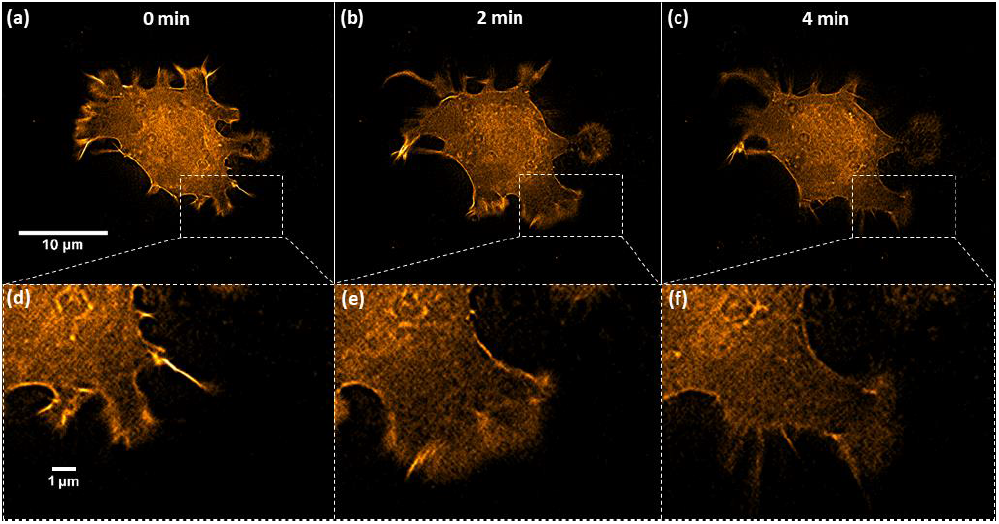
Live-cell time-lapse images of fluorescent actin in a MC3T3-E1 cell at **(a,d)** 0 min, **(b,e)** 2 mins, and **(c,f)** 4 mins. **(d-f)** represent higher magnification images of the highlighted regions in **(a-c)**. Transparent TiO2 NTs are 56 ± 6 nm in diameter, 633 ± 47 nm long, and cells were seeded 20 mins prior to imaging.

## 4. Conclusions

This work began by identifying and establishing the relationship of TiO_2_ NT fabrication conditions to produce robust TiO_2_ NT coatings with tailorable physical characteristics (i.e. length and diameter). Success in this task was demonstrated by production of stable TiO_2_ NT coatings with tailorable diameters (OD of ∼56-116 nm), wall thicknesses (∼12-28 nm), and lengths (∼0.6-6.4 μm) made possible by optimized TiO2 NT fabrication techniques. These techniques were translated to produce transparent TiO_2_ NTs from physical vapor deposited (PVD) Ti. All TiO_2_ NT coatings produced from these methods were found to have high optical transparencies (≥60%). Epifluorescence microscopy of fluorescently-tagged vinculin and f-actin in live pre-osteoblast (MC3T3-E1) cells confirmed suitability of the transparent TiO_2_ NT platform for use in future experiments focused on investigating dynamic responses of cells interacting with nanostructured substrates in real-time. These tailorable, transparent TiO_2_ NT platforms can be used in conjunction with standard and advanced fluorescence microscopy techniques to identify, quantify, and characterize dynamic cellular processes, and may be amenable to a variety of advanced imaging techniques employed to elucidate the mechanisms by which nanostructure influences cell behavior.

## 5. Acknowledgments

This material is based upon work supported by the National Science Foundation/EPSCoR Cooperative Agreement #IIA-1355423 and by the State of South Dakota. Any opinions, findings, and conclusions or recommendations expressed in this material are those of the author(s) and do not necessarily reflect the views of the National Science Foundation. Thanks to Dr. Frank Kustas and Jacob Petersen for assistance with physical vapor deposition and atomic force microscopy, respectfully.

## 7. Data Availability

1. The raw/processed data required to reproduce these findings cannot be shared at this time as the data also forms part of an ongoing study.

